# Identification of secreted proteins by comparison of protein abundance in conditioned media and cell lysates

**DOI:** 10.1101/2022.06.16.496407

**Authors:** Prabhodh S. Abbineni, Vi T. Tang, Felipe da Veiga Leprevost, Venkatesha Basrur, Jie Xiang, Alexey I. Nesvizhskii, David Ginsburg

## Abstract

Analysis of the full spectrum of secreted proteins in cell culture is complicated by leakage of intracellular proteins from damaged cells. To address this issue, we compared the abundance of individual proteins between the cell lysate and the conditioned medium, reasoning that secreted proteins should be relatively more abundant in the conditioned medium. Marked enrichment for signal-peptide-bearing proteins with increasing conditioned media to cell lysate ratio, as well loss of this signal following brefeldin A treatment, confirmed the sensitivity and specificity of this approach. The subset of proteins demonstrating increased conditioned media to cell lysate ratio in the presence of Brefeldin A identified candidates for unconventional secretion via a pathway independent of ER to Golgi trafficking.

## Introduction

Approximately 34% of genes in the mammalian genome encode proteins destined for the secretory pathway, with these secreted proteins serving a variety of physiological roles [1, 2]. Proteins bearing signal-peptides and/or transmembrane domains are co-translationally inserted into the endoplasmic reticulum (ER) where they are folded prior to undergoing anterograde transport to the Golgi *via* coatomer protein complex II (COPII) coated vesicles or tubular structures [3-5]. In the Golgi, these proteins are then packaged into mature vesicles that enter constitutive or regulated exocytosis pathways for delivery to various intracellular compartments, or fusion with the plasma membrane for secretion into the extracellular space or insertion into the plasma membrane. While the majority of secreted proteins utilize the ER-Golgi-Secretory vesicle pathway, a subset of cytoplasmic proteins have been shown to be secreted *via* ER-Golgi independent “unconventional” secretory routes [6-8]. Following synthesis in the cytoplasm, unconventionally secreted proteins, which include some cytokines and angiogenic growth factors, are secreted either *via* protein conducting pores that form in the plasma membrane, or *via* vesicular intermediates derived from the auto/endo-lysosomal system[8-10].

Characterizing the entire set of secreted proteins, i.e. the secretome, using cell culture-based model systems has been a long standing challenge [2]. Several studies have focused on identifying changes to the secretome under pathophysiological conditions to identify biomarkers for disease, or following inhibition of specific components of the secretory pathway with the aim of identifying specific regulators of select secreted proteins [11-14]. However, a major challenge in studying the secretome of cultured cells is contamination by intracellular proteins released into the media due to cellular injury, which confounds identification of bona-fide secreted proteins [15]. To circumvent this issue, we compared the abundance of proteins identified in the cell lysate and conditioned media. We reasoned that secreted proteins should be more abundant in the media relative to the cell lysates, whereas intracellular proteins that are released into the media due to cell injury should be present at a lower ratio in the media compared to the cell lysates. Previous studies have utilized a similar approach using label-free quantification-based mass spectrometry whereby spectral counts from conditioned media and cell lysates are compared to identify secreted proteins, or have applied isobaric tags for relative and absolute quantitation (ITRAQ) reagents for label-based quantification of media and lysate proteins [16-18]. In order to independently test this approach, we utilized isobaric tandem-mass-tags (TMT) to label proteins in the cell lysate and conditioned media of Huh7 hepatoma cells, followed by high throughput mass spectrometry to compare the abundance of proteins in conditioned media relative to cell lysates. This approach distinguished between intracellular and secreted proteins with high specificity. Treatment with brefeldin-A (BFA), an inhibitor of ER-Golgi trafficking, provided additional validation of this approach and identified candidate proteins trafficked *via* BFA-resistant unconventional secretion pathways. The approach described here is readily applicable to characterizing secretomes from a wide range of cell types.

## Methods

### Cell culture, and BFA treatment

Huh7 cells [19] were cultured in DMEM Glutamax media (ThermoFisher Scientific, Waltham MA, 10569-044) supplemented with 10% fetal bovine serum (MilliporeSigma, Burlington MA, F8067) and penicillin/streptomycin (ThermoFisher Scientific, Waltham MA, 15140-122). Cells were maintained at 20-85% confluence and passaged every 3-4 days. For BFA experiments, Huh7 cells were treated with 1 µg/ml of BFA (BioLegend, San Diego CA, 420601) and after incubation for 4 hours in serum-free, phenol red-free DMEM, conditioned medium and cell lysate were collected and processed as described below.

### Conditioned medium collection

Conditioned media were centrifuged at 2500g at 4ºC for 20 minutes to pellet cell debris, followed by ultracentrifugation at 120,000g at 4ºC for 90 minutes to pellet exosomes [20]. Following ultracentrifugation, the supernatants were concentrated using a 3 kDa molecular weight cutoff concentrator (MilliporeSigma, Burlington MA, UFC900324). Protein concentrations were determined by DC protein assay (Bio-Rad, Hercules CA, 500-011).

### Cell lysate collection

Following removal of conditioned media, cells were rinsed twice in cold (4ºC) PBS, followed by the addition of 2 ml of RIPA buffer (Thermo Scientific, catalog # 89900) containing a protease inhibitor cocktail (cOmplete™, Mini Protease Inhibitor Cocktail, catalog # 11836153001). Cells were then detached using a cell scraper and collected into 15 ml conical tubes. Cell suspensions were sonicated and rotated end-over-end for 1 hour. Next, lysates were centrifuged at 21,000g at 4ºC for 45 minutes. The supernatants were then transferred to a new Eppendorf tube and the protein concentrations determined by DC protein assay).

### Immunoblotting

Cell lysates were diluted to 2 µg/µl in Laemmli buffer (Bio-Rad, Hercules CA, 1610747) and heated at 95ºC for 5 minutes and resolved using a 4-20% tris-glycine gel (ThermoFisher Scientific, Waltham MA, XP04200). Next, samples were transferred to nitrocellulose membranes (ThermoFisher Scientific, Waltham MA, LC2000) and probed with antibodies against PCSK9 (abcam, Cambridge UK, ab181142, 1:1000), and ß-actin (Santa Cruz Biotechnology, Dallas TX, sc-47778, 1:1000). Chemiluminescence-based quantification of PCSK9 abundance in Huh7 cell lysates and conditioned media following immunoblotting was performed using the Azure c600 imager (Azure Biosystems, CA, USA), and ImageJ for image analysis [21].

### Mass spectrometry

Labeling of peptides with TMT reagents followed by TMT proteomics was performed as previously described [22]. 75 µg lysate and 50 µg media samples were proteolysed and labeled with TMT 6-plex according to the manufacturer’s protocol (ThermoFisher). Briefly, upon reduction (5 mM DTT, for 30 min at 45 C) and alkylation (15 mM 2-chloroacetamide, for 30 min at room temperature) of cysteines, the proteins were precipitated by adding 6 volumes of ice cold acetone followed by overnight incubation at -20° C. The precipitate was collected by centrifugation, and the pellet was allowed to air dry. The pellet was resuspended in 0.1M Triethylammonium bicarbonate (TEAB) and overnight (∼16 h) digestion was performed with trypsin/Lys-C mix (1:25 protease:protein; Promega) at 37° C with constant mixing using a thermomixer. The TMT 6-plex reagents were dissolved in 41 µl of anhydrous acetonitrile and labeling was performed by transferring the entire digest to TMT reagent vial and incubating at room temperature for 1 h. Reaction was quenched by adding 8 µl of 5% hydroxyl amine and further 15 min incubation. Labeled samples were mixed together, and dried using a vacufuge. An offline fractionation of the combined sample (∼200 µg) into 8 fractions was performed using high pH reversed-phase peptide fractionation kit according to the manufacturer’s protocol (Pierce; Cat #84868). Fractions were dried and reconstituted in 9 µl of 0.1% formic acid/2% acetonitrile in preparation for LC-MS/MS analysis.

### Liquid chromatography-mass spectrometry analysis (LC-multinotch MS3)

In order to obtain superior quantitation accuracy, we employed multinotch-MS3 (McAlister GC, ref below) which minimizes the reporter ion ratio distortion resulting from fragmentation of co-isolated peptides during MS analysis [23]. Orbitrap Fusion (Thermo Fisher Scientific) and RSLC Ultimate 3000 nano-UPLC (Dionex) was used to acquire the data. Two µl of the sample was resolved on a PepMap RSLC C18 column (75 µm i.d. x 50 cm; Thermo Scientific) at the flow-rate of 300 nl/min using 0.1% formic acid/acetonitrile gradient system (2-22% acetonitrile in 150 min; 22-32% acetonitrile in 40 min; 20 min wash at 90% followed by 50 min re-equilibration) and directly spray onto the mass spectrometer using EasySpray source (Thermo Fisher Scientific). Mass spectrometer was set to collect one MS1 scan (Orbitrap; 120K resolution; AGC target 2×10^5^; max IT 100 ms) followed by data-dependent, “Top Speed” (3 seconds) MS2 scans (collision induced dissociation; ion trap; NCE 35; AGC 5×10^3^; max IT 100 ms). For multinotch-MS3, top 10 precursors from each MS2 were fragmented by HCD followed by Orbitrap analysis (NCE 55; 60K resolution; AGC 5×10^4^; max IT 120 ms, 100-500 m/z scan range).

### Protein identification and quantification

Raw mass spectrometry files were converted into open mzML format using the msconvert utility of the Proteowizard software suite, and analyzed using FragPipe computational platform (https://fragpipe.nesvilab.org/) using default TMT10-MS3 workflow. MS/MS spectra were searched using the database search tool MSFragger v3.4 [24, 25] against an *Homo sapiens* UniprotKB/SwissProt protein sequence database appended with an equal number of decoy sequences (downloaded on January 31, 2022). Whole cell lysate MS/MS spectra were searched using a precursor-ion mass tolerance of 20 ppm, and allowing C12/C13 isotope errors 0/1/2/3. Mass calibration and parameter optimization were enabled. Cysteine carbamidomethylation (+57.0215), and lysine TMT labeling (+229.1629) were specified as fixed modifications, and methionine oxidation (+15.9949), N-terminal acetylation (+42.01060), and TMT labeling (+229.1629) of peptide N terminus and serine residues were specified as variable modifications. The search was restricted to tryptic peptides, allowing up to two missed cleavage sites. Peptide to spectrum matches (PSMs) were further processed using Percolator [26] to compute the posterior error probability, which was then converted to posterior probability of correct identification for each PSM. The resulting files from Percolator were converted to pep.xml format, and pep.xml files from both TMT experiments (the secretome and the whole lysate experiment) were then processed together to assemble peptides into proteins (protein inference) using ProteinProphet [27] run via the Philosopher toolkit v4.2.1 [28] to create a combined file (in prot.xml format) of high confidence protein groups. The combined prot.xml file and the individual PSM lists for each of the two TMT experiment were further processed using the Philosopher filter command as follows. Each peptide was assigned either as a unique peptide to a particular protein group or assigned as a razor peptide to a single protein group that had the most peptide evidence. The protein groups assembled by ProteinProphet were filtered to 1% protein-level False Discovery Rate (FDR) using the target-decoy strategy and the best peptide approach (allowing both unique and razor peptides). The PSM lists were filtered using a sequential FDR strategy, retaining only those PSMs that passed 1% PSM-level FDR filter and mapped to proteins that also passed the global 1% protein-level FDR filter. In addition, for all PSMs corresponding to a TMT-labeled peptide, TMT reporter ion intensities were extracted from the MS/MS scans (using 0.002 Da window) and the precursor ion purity scores were calculated using the intensity of the sequenced precursor ion and that of other interfering ions observed in MS1 data (within a 0.7 Da isolation window). The PSM output files were further processed using TMT-Integrator v3.2.1 to generate summary reports at the protein level. TMT-Integrator [29] used as input the PSM tables generated by the Philosopher pipeline as described above and created integrated reports with quantification across all samples. First, PSMs were filtered to remove all entries that did not pass at least one of the quality filters, such as PSMs with (a) no TMT label; (b) precursor-ion purity less than 50%; (c) summed reporter ion intensity (across all channels) in the lower 5% percentile of all PSMs in the corresponding PSM.tsv file. In the case of redundant PSMs (i.e., multiple PSMs in the same MS run sample corresponding the same peptide ion), only the single PSM with the highest summed TMT intensity was retained for subsequent analysis. Both unique and razor peptides were used for quantification, while PSMs mapping to common external contaminant proteins (that were included in the searched protein sequence database) were excluded. Next, for each PSM the intensity in each TMT channel was converted into log2-based ratio to the reference using the virtual reference approach of TMT-Integrator. The PSMs were grouped to the protein level, and the protein ratios were computed as the median of the corresponding PSM ratios after outlier removal. Protein ratios were then converted back to absolute protein intensity in each sample by using the reference protein intensity estimated (separately for each experiment, the secretome and the whole lysate), using the sum of all MS2 reporter ions from all corresponding PSMs.

In order to calculate media/lysate abundance ratio, we used the absolute protein intensities values with log2(media/lysate) ratio as input. Statistical analysis of changes to protein abundance and media/lysate ratio following BFA treatment relative to untreated controls was performed using the limma statistical package [30].

## Results

### Identification of secreted proteins by comparison of protein abundance in the cell lysate and conditioned media

We reasoned that proteins secreted by cultured cells should exhibit significantly greater abundance in the conditioned media relative to the cell lysate, when compared to non-secreted proteins. For this analysis, protein preparations from conditioned media and cell lysates of Huh7 cells were subjected to quantitative proteomic analysis using TMT and LC-MS/MS. Samples from different conditions were pooled after TMT labelling, with the lysates and media fractions analyzed in independent MS runs (**Fig. 1A**).

**Figure 1.**
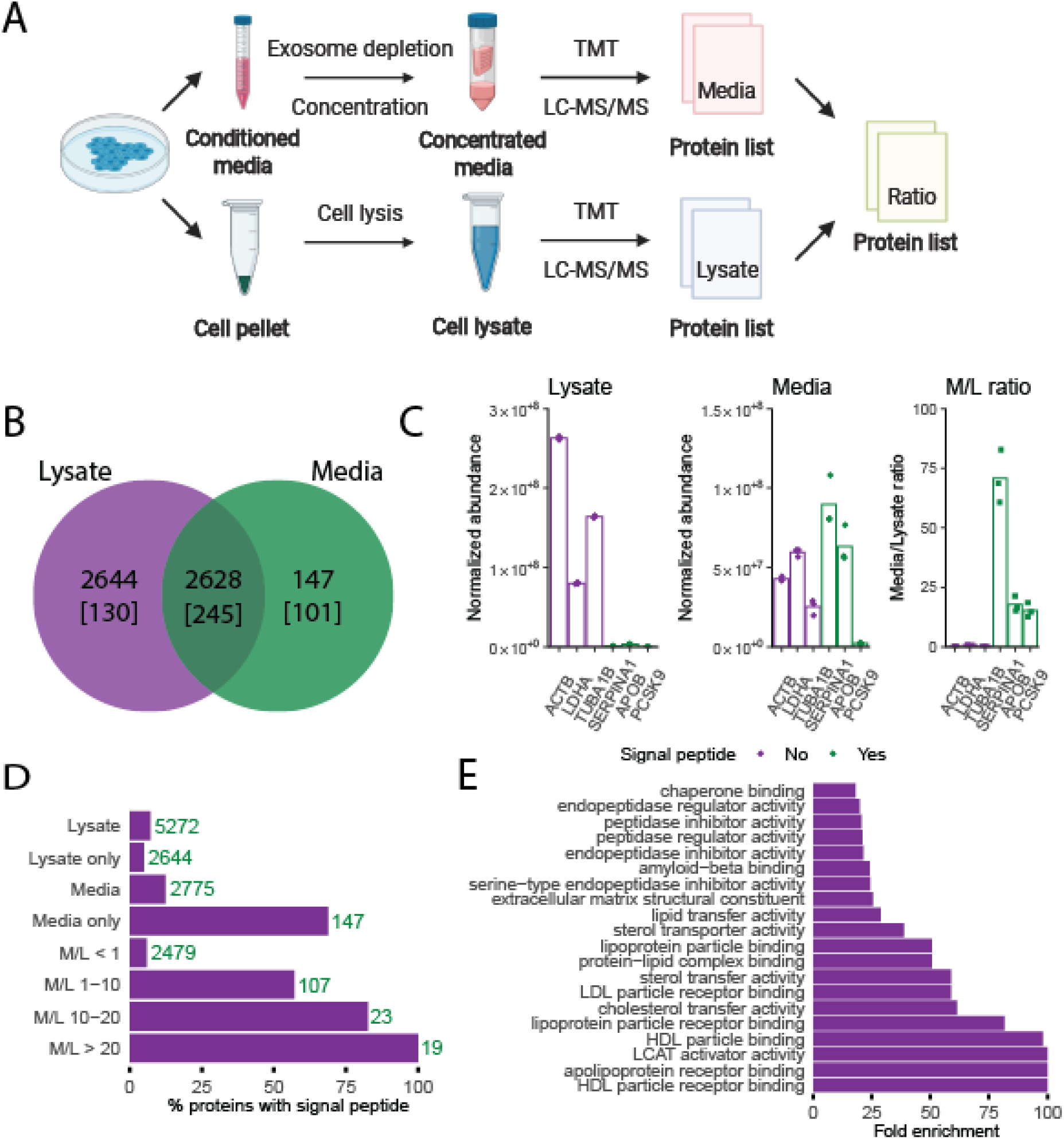
Comparison of protein abundance in the cell lysate and conditioned media enables identification of secreted proteins. **(A)** Experimental approach: Proteins isolated from cell lysates and conditioned media were labelled with tandem-mass-tag (TMT) reagents prior to LC-MS/MS analysis to quantify relative protein abundance. A media/lysate (M/L) abundance ratio was calculated for all proteins identified in both fractions. **(B)** Venn diagram showing the overlap in proteins detected in the cell lysates and conditioned media. The number of proteins with signal peptides present in each fraction is indicated in brackets. **(C)** The abundance of known cytoplasmic proteins (ACTB, LDHA, TUBA1B) and well documented secreted proteins (SERPINB1, APOB, PCSK9) in the lysate and media fraction, and their M/L ratio. **(D)** Proteins were binned into discreet M/L ratio intervals, and the percentage of proteins with signal peptides in each bin was calculated. The number of proteins present in each bin is indicated. **(E)** Gene ontology (GO) analysis for molecular functions for the set of proteins with M/L ratio greater than 10.

A total of 5419 unique proteins were identified in the lysate and media fractions across all three independent replicates, with 48% of these proteins identified in both the media and lysate fractions (**Fig. 1B**). Only 12.5% of the proteins detected in the media contained signal-peptides (**Fig. 1B**), a 1.75-fold enrichment relative to the cell lysate. These data suggest that a large subset of the proteins detected in the media represent cytoplasmic/intracellular protein contamination, likely due to cell damage or lysis. We then calculated the media to lysate abundance ratio (M/L ratio) for 3 known cytoplasmic proteins (all lacking signal peptides) that were detected in the media, actin beta (ACTB), lactate dehydrogenase A (LDHA) and tubulin alpha-1B chain (TUBA1B), and 3 well documented signal peptide-containing, liver-specific secreted proteins, alpha-1-antitrypsin (SERPINA1), apolipoprotein B (APOB), and proprotein convertase subtilisin/kexin type 9 (PCKS9) (**Fig. 1C**). As expected, all 3 signal peptide containing proteins exhibit significantly higher M/L ratios (∼12-80, **Fig. 1C M/L ratio**) relative to the intracellular proteins (∼0.1-0.8, *p* = 0.0019). Calculation of media to lysate abundance ratios (M/L ratio) for all 2628 proteins identified in both the media and lysate fractions is shown in **Fig. 1D**. The fraction of proteins with signal peptides steadily increased with greater M/L ratio, consistent with greater enrichment of secreted proteins in the media relative to the lysate, and validating the media/lysate ratio analysis approach.

Gene ontology analysis of proteins with M/L ratios greater than 10 revealed significant enrichment for proteins involved in multiple lipoprotein and cholesterol metabolism processes (**Fig. 1E**), consistent with the hepatocyte origin of the Huh7 cell line. The majority, 71.3%, of proteins detected solely in the media contained signal peptides (101 out of 147), with this subset largely composed of low abundance secreted proteins, likely explaining their failure to be detected in the lysate fraction. The remaining proteins without signal peptides represent potential unconventionally secreted proteins (see below). Only ∼5% of proteins detected solely in the lysate contained signal peptides, which are largely intracellular membrane associated proteins with transmembrane domains (91 out of 130 proteins), or soluble proteins contained within the lumen of intracellular organelles, including the lysosomal proteins, cathepsin L2 and galactocerebrosidase.

### BFA inhibits ER-Golgi trafficking and further validates the M/L ratio analysis approach

We next treated Huh7 cells with BFA, an Arf1 GTPase inhibitor that blocks ER-Golgi transport and conventional protein secretion (**Fig. 2A**). Western blotting for the conventionally secreted protein, PCSK9, with or without BFA treatment is shown in **Fig. 2B**. As expected, PCSK9 accumulation in the cell lysate and depletion from the conditioned media is observed following BFA treatment (**Fig. 2C**). Proteomic analysis of control and BFA treated cell lysates and conditioned media by TMT LC-MS/MS analysis is shown in **Fig. 2D-F**.

**Figure 2.**
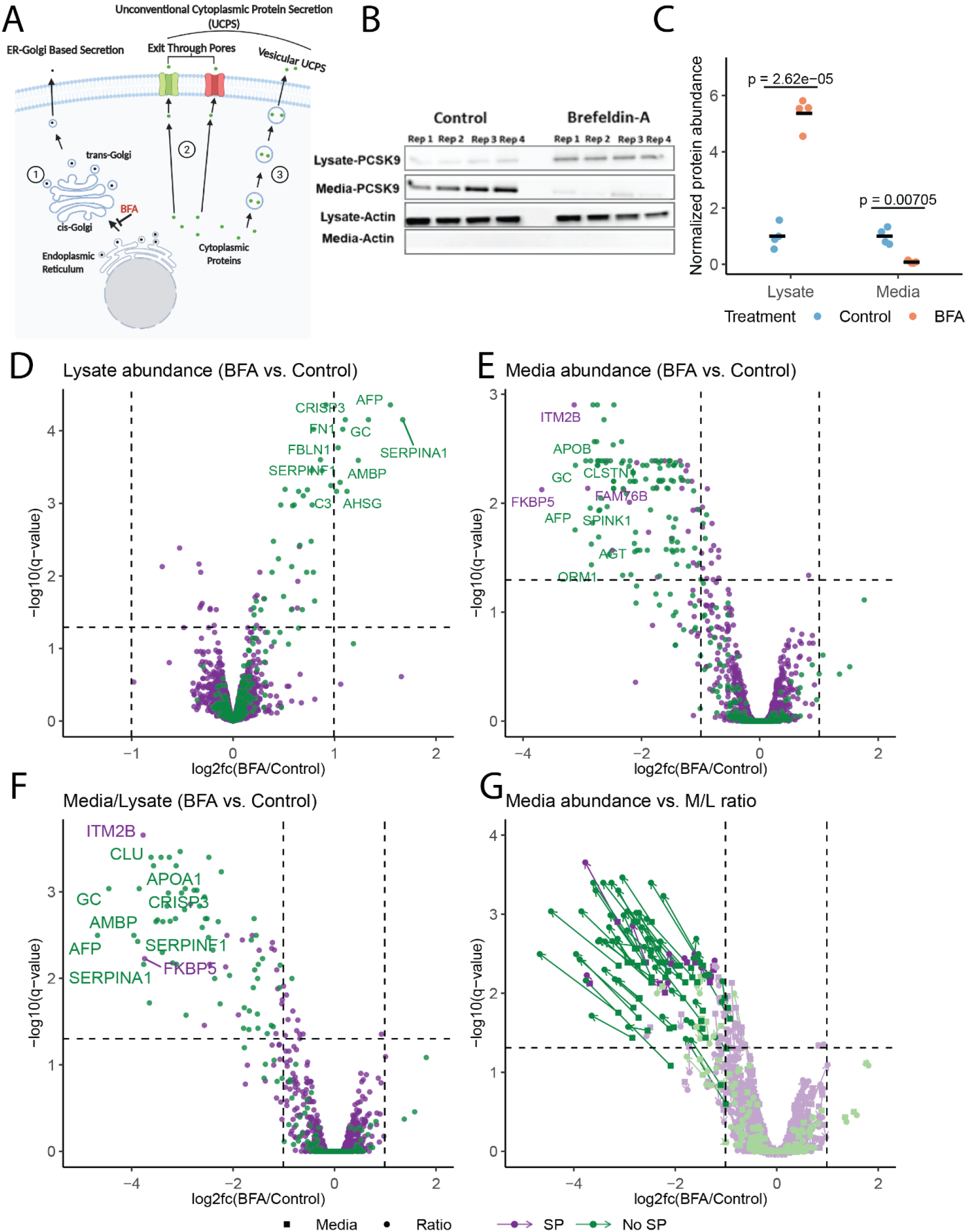
M/L ratio analysis combined with BFA treatment enables high-sensitivity identification of conventionally secreted proteins. **(A)** Proteins bearing signal peptides are co-translationally inserted into the ER and utilize the conventional ER-Golgi-Secretory vesicle/tubule route (pathway (1)) to exit the cell or to reach other intracellular destinations. The process of ER-Golgi transport is blocked by BFA. A number of cytoplasmic proteins are known to be secreted unconventionally *via* ER-Golgi independent secretory routes by either translocating through plasma membrane pores that are formed by the secreted protein itself, or by other pore-forming proteins (pathway(2)), or by entering membrane bound organelles that fuse with the plasma membrane (pathway (3)). These latter 2 groups of proteins should be resistant to BFA. **(B)** Immunoblot analysis of PCSK9 abundance in the Huh7 cell lysate and conditioned media following treatment with 1 ug/ml BFA. Four biological replicates are labeled Rep1-4. **(C)** Chemiluminescence-based quantification of PCSK9 abundance in Huh7 cell lysates and conditioned media following immunoblotting. BFA treated samples were normalized to controls. (**D-G**) Volcano plots [43] showing changes to the abundance of proteins in the cell lysate (**D**) and conditioned media **(E)**, and changes to M/L ratio **(F)** following BFA treatment measured by TMT-based mass spectrometry. The l2fc and statistical significance are plotted on the x and y-axis, respectively. Signal peptide containing proteins are indicated in green, and non-signal peptide containing in purple. (**G**) Comparison of l2fc following BFA treatment for individual proteins comparing protein abundance in the media alone (squares) to the M/L ratio (circles).

In **Fig. 2 D and E**, the volcano plots show the log2 fold changes (l2fc) in the abundance of proteins in the cell lysate and conditioned media, respectively, following BFA treatment. A positive l2fc indicates increased abundance in response to BFA treatment, and a negative l2fc indicates decreased abundance. Following BFA treatment, most signal peptide containing proteins exhibit positive l2fc in the lysate (**Fig. 2D**), and negative l2fc in the media (**Fig. 2E**), consistent with BFA inhibition of conventional protein secretion resulting in depletion of secreted proteins from the conditioned media and accumulation in the cell lysate. Signal peptide containing proteins that don’t change in abundance following BFA treatment are comprised primarily of either membrane proteins (38.6%, 93 out of 241 proteins), or are confined to the lumen of intracellular compartments as indicated by GO cellular component analysis. In order to account for protein abundance changes in both the lysate and media fractions as described above, we calculated l2fc of M/L ratios following BFA treatment (**Fig. 2F**). A negative l2fc indicates a decrease in M/L in response to BFA treatment (depletion from the media and/or accumulation in the lysate). 75% of proteins with negative l2fc contain signal peptides, further confirming the robustness of this analysis approach. **Fig. 2G** compares the l2fc observed for protein abundance in the conditioned media to the l2fc for the M/L ratio. The M/L ratio analysis, which considers protein abundance in both the media and lysate fractions, provides a higher sensitivity for the identification of secreted proteins, with considerably larger (negative) l2fc compared to analysis of the conditioned media alone for the same protein. The 25% of proteins sensitive to BFA treatment but lacking signal peptides are limited to membrane proteins (e.g, integral membrane protein 2B (ITM2B)) with transmembrane domains and/or signal anchors that target them to the endomembrane system, or soluble proteins that potentially have cryptic internal signal peptides that target them to the ER (e.g, FKBP prolyl isomerase 5 (FKBP5)). The majority of proteins detected solely in the media contain signal peptides (71%), with most of these depleted from the media upon BFA treatment (**Fig. S1 A**).

### Identification of unconventionally secreted proteins

All proteins with an M/L ratio greater than 20 (19 proteins) contain a signal peptide, suggesting that the most efficiently secreted proteins in this cell type use the conventional ER-Golgi secretory pathway. In contrast, 17% of proteins with M/L ratios of 10 – 20 (4 out of 23 proteins) and 43% of proteins with M/L ratios of 1 – 10 (46 out of 107 proteins) lack a signal-peptide, suggesting that a subset of these proteins may be secreted via unconventional, ER-Golgi independent pathways (**Fig. 1D)**. Of the 10 proteins without signal peptides with the highest M/L ratios, 9 are resistant to BFA treatment (**Fig. 3A-B**). Among this group, the protein with the highest M/L ratio, metallothionein 2A (MT2A), has previously been shown to be secreted via ER-Golgi independent mechanisms [31, 32], and it’s identification in our dataset further confirms the utility of our approach to identify both conventionally and unconventionally secreted proteins. The 1 exception among the top 10 proteins without signal peptides with the highest M/L ratios that was BFA sensitive, FKBP5, is likely still dependent on the conventional secretory pathway, despite the absence of an annotated signal peptide. 2 of the 10 proteins, inositol 1,4,5-triphosphate receptor associated 2 (IRAG2) and golgi membrane protein 1 (GOLM1), contain transmembrane domains that likely traffic them to the ER, but their resistance to BFA indicates that they may be stored in a post-Golgi compartment for secretion, as BFA inhibits ER-Golgi trafficking. The remaining 6 proteins in this list (gene names: LZTS1, ZBTB8OS, ANKRD40, FNTA, HECTD3, and GFUS) are resistant to BFA treatment (**Fig. 3B**) and lack signal peptides and transmembrane domains. Although all have been reported to have intracellular roles in regulating mitosis [33], transcription [34], and posttranslational modifications [35-37], and are not known to be secreted, our findings raise the possibility that one or more of these proteins may indeed be secreted *via* UCPS with an as yet unknown extracellular function.

**Figure 3.**
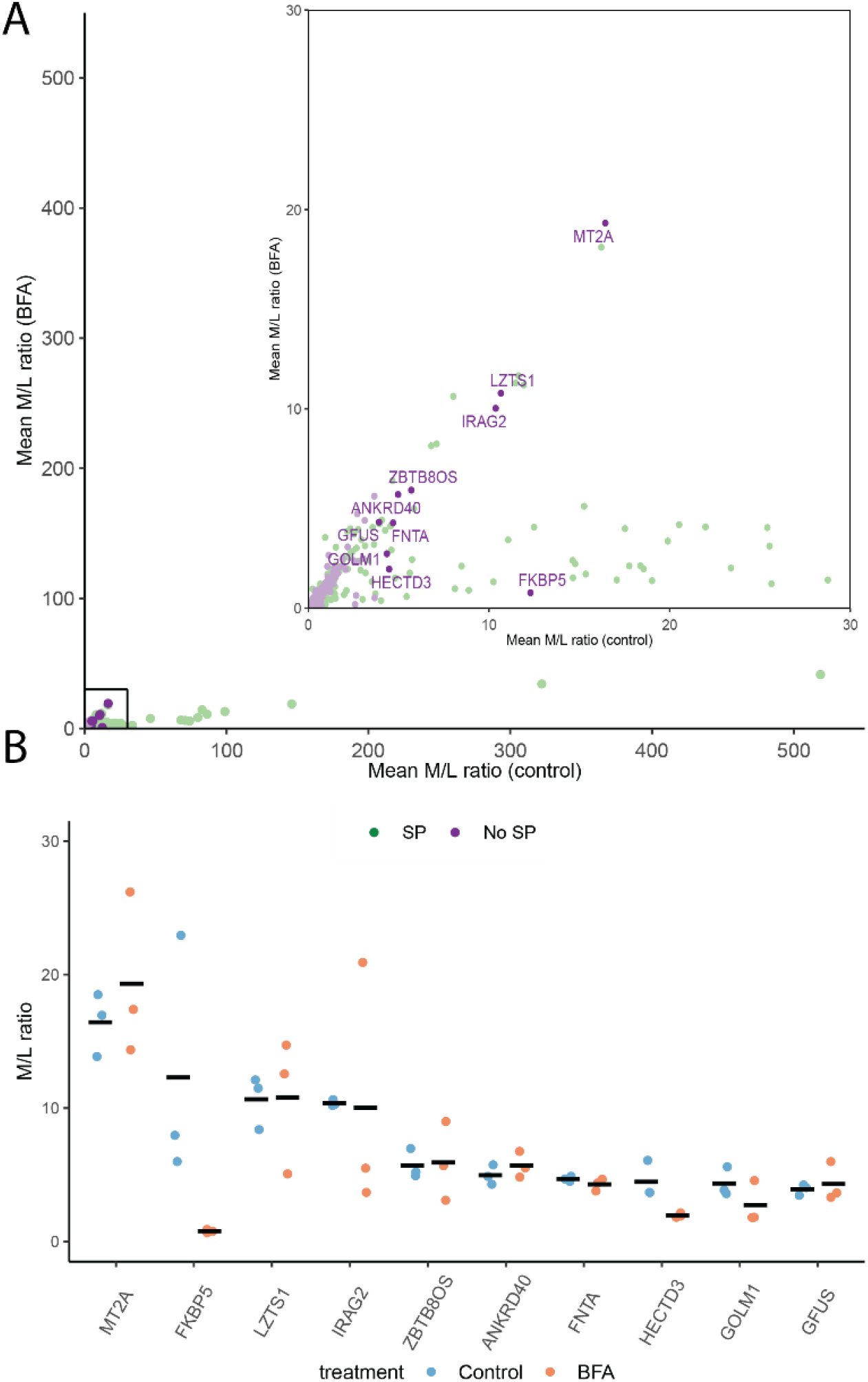
Candidate proteins secreted *via* unconventional secretory pathways. **(A)** M/L ratio of control and BFA treated cells are plotted on the x and y-axis, respectively. The inset shows proteins with a M/L ratio ranging from 0-30. Proteins sensitive to BFA have lower M/L ratios relative to untreated controls. The top 10 signal-peptide lacking proteins with the highest M/L ratios (candidate unconventionally secreted proteins) are identified by gene name. **(B)** M/L ratio of candidate unconventionally secreted proteins and sensitivity to BFA.

## Discussion

Secretion of proteins into the extracellular space is a fundamental process that occurs in eukaryotic cells. In multicellular organisms, secreted proteins play several essential roles, including intercellular communication, maintaining metabolic homeostasis, immune reactions, and neurotransmission. Efforts to define the set of proteins that are secreted, called the secretome, has faced significant obstacles. Currently, the human secretome is defined primarily based on computational predictions of proteins that have a signal peptide and exclusion of transmembrane proteins [1, 2]. This approach has several shortcomings, including inaccuracy of algorithms for identification of proteins with a signal peptide or a transmembrane domain and omission of unconventionally secreted proteins. There are also significant challenges in defining a secretome experimentally. In vivo studies to identify circulating proteins in the plasma by mass spectrometry faces the problem of variable protein abundance and the difficulty of detecting low abundance proteins [38]. This approach also excludes the large number of secreted proteins that function locally and do not enter the circulation. In vitro studies of cultured cells circumvent some of these problems and also allow selection of a specific cell type of interest. However, this approach is subject to a high false positive rate due to contamination by intracellular proteins that leak into the conditioned media from damaged cells.

Here, we propose an alternative approach that facilitates efficient and accurate identification of secreted proteins by analysis of both conditioned media and cell lysates. We show that in contrast to intracellular proteins, authentically secreted proteins are significantly more abundant in the conditioned media relative to the cell lysate. By analyzing an M/L ratio for each protein, we were able to clearly distinguish secreted proteins from those that typically remain intracellular. The efficacy of this ratiometric approach to identify secreted proteins was further confirmed by our observation that treatment with BFA significantly decrease the M/L ratio of proteins with a signal peptide compared to those without. Furthermore, as this approach does not require genetic manipulation or metabolic labeling, it can be widely applied to both primary and immortalized cell culture systems.

In contrast to proteins traversing the conventional ER to Golgi secretory pathway, no definitive structural features analogous to the signal peptide have been identified that target cytoplasmic proteins for UCPS. Although algorithms such as SecretomeP have been designed to predict unconventionally secreted proteins based on shared physiochemical features (isoelectric point, number of charged residues, secondary structure, etc.) (7), a recent analysis of SecretomeP and similar programs tested for known unconventionally secreted and non-secreted proteins demonstrated low sensitivity and a high false positive rate (8). Thus, while hundreds of proteins have been proposed to use unconventional secretory routes, few have been convincingly validated, and no standard methods are available to detect unconventionally secreted proteins. Our ratiometric analysis approach combined with the use of BFA to block ER-Golgi trafficking facilitated the identification of candidate unconventionally secreted cargoes. This method is also readily adaptable to identify unconventionally secreted proteins from diverse cell lines or cultured primary cells.

Stühler and colleagues recently pioneered the approach of comparing protein abundance in the cellular lysate fraction and conditioned media to identify secreted proteins [15, 16, 39]. We independently confirm the utility of this approach by using TMT-based labeling of lysate and media proteins, followed by label-based quantification of TMT data. Previous analyses have classified proteins as secreted if their mean abundance in the conditioned media was at least 1.5-fold higher relative to the cell lysate [17]. To identify candidate UCPS cargoes, we applied more stringent criteria based on prioritizing signal-peptide lacking, BFA-resistant proteins with the highest M/L ratios. Our observation of a strong concordance between increasing M/L ratio and higher fractions of signal peptide containing proteins suggests that the UCPS candidates with the highest M/L ratios are most likely to represent true positives. Future studies should uncover extracellular roles for these novel UCPS cargoes and characterize molecular mechanisms that regulate their release from the cell.

The M/L ratio approach relies on the assumption that secreted proteins will be more abundant in the media relative to the lysate; thus, proteins that have dual intra-and extracellular roles that are present in similar quantities in the media and lysate may fail to be identified by this approach. Examples of such proteins include the unconventionally secreted proteins fibroblast growth factor 2 [40], galectin-3 [41], and histone-3A [42]. Furthermore, as this analysis is limited to proteins detected in the conditioned media, secreted proteins that reside on the cell surface or remain insoluble in the subcellular matrix will also be missed by this approach. Thus, the introduction of additional biochemical procedures to isolate these proteins will be required to gain a more comprehensive view of the full secretome.

**Figure S1.**
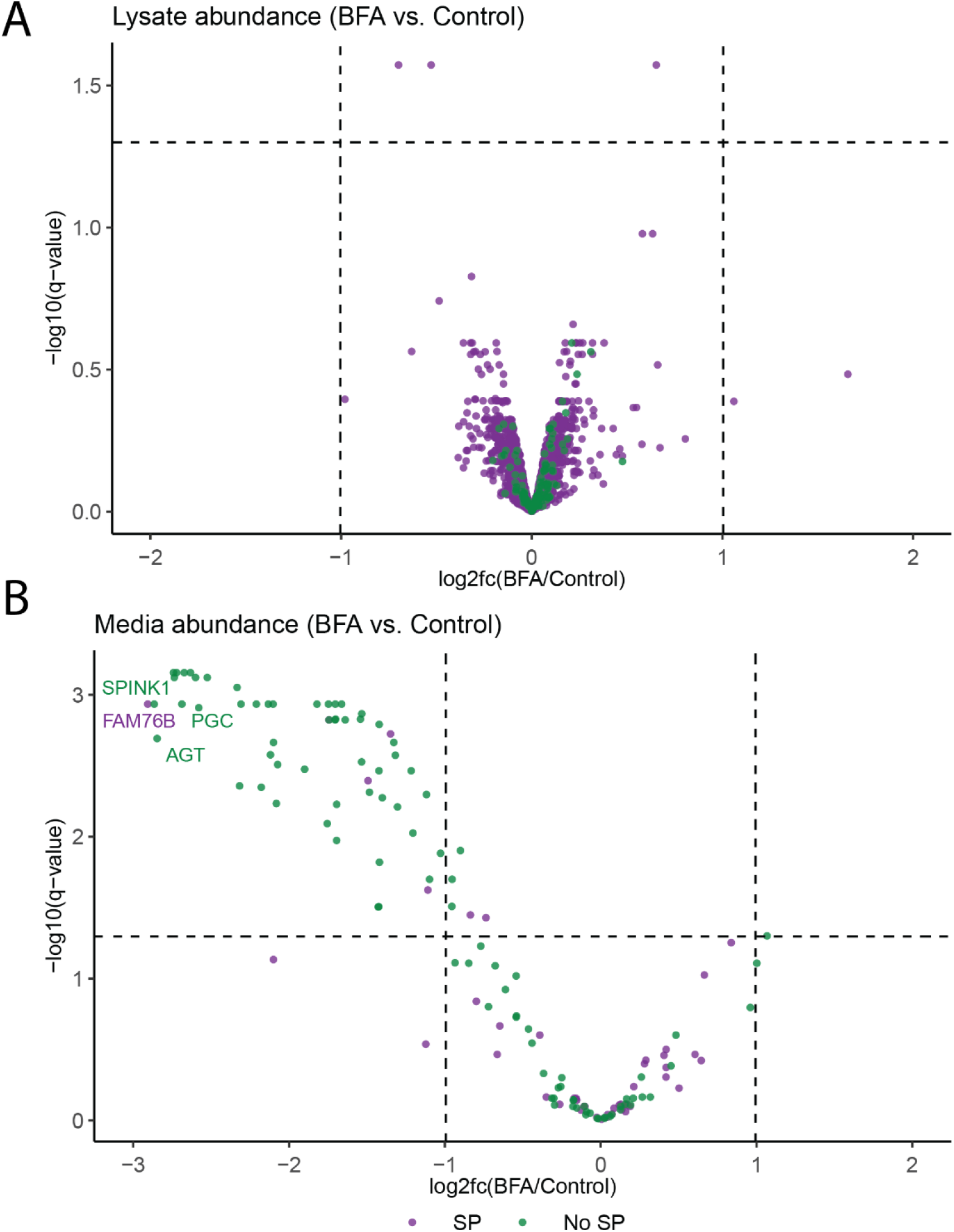
Proteins detected solely in the media fraction are sensitive to BFA. Volcano plots showing changes to the abundance of proteins detected solely in the cell lysate (**A**) and conditioned media **(B)** following BFA treatment measured by TMT-based mass spectrometry.

## Notes

### Competing Interest Statement

The authors have declared no competing interest.

